# Collective feature selection to identify crucial epistatic variants

**DOI:** 10.1101/293365

**Authors:** Shefali S. Verma, Anastasia Lucas, Xinyuan Zhang, Yogasudha Veturi, Scott Dudek, Binglan Li, Ruowang Li, Ryan Urbanowicz, Jason H. Moore, Dokyoon Kim, Marylyn D. Ritchie

## Abstract

**Background:** Machine learning methods have gained popularity and practicality in identifying linear and non-linear effects of variants associated with complex disease/traits. Detection of epistatic interactions still remains a challenge due to the large number of features and relatively small sample size as input, thus leading to the so-called “short fat data” problem. The efficiency of machine learning methods can be increased by limiting the number of input features. Thus, it is very important to perform variable selection before searching for epistasis. Many methods have been evaluated and proposed to perform feature selection, but no single method works best in all scenarios. We demonstrate this by conducting two separate simulation analyses to evaluate the proposed collective feature selection approach.

**Results:** Through our simulation study we propose a collective feature selection approach to select features that are in the “union” of the best performing methods. We explored various parametric, non-parametric, and data mining approaches to perform feature selection. We choose our top performing methods to select the union of the resulting variables based on a user-defined percentage of variants selected from each method to take to downstream analysis. Our simulation analysis shows that non-parametric data mining approaches, such as MDR, may work best under one simulation criteria for the high effect size (penetrance) datasets, while non-parametric methods designed for feature selection, such as Ranger and Gradient boosting, work best under other simulation criteria. Thus, using a collective approach proves to be more beneficial for selecting variables with epistatic effects also in low effect size datasets and different genetic architectures. Following this, we applied our proposed collective feature selection approach to select the top 1% of variables to identify potential interacting variables associated with Body Mass Index (BMI) in ~44,000 samples obtained from Geisinger’s MyCode Community Health Initiative (on behalf of DiscovEHR collaboration).

**Conclusions:** In this study, we were able to show that selecting variables using a collective feature selection approach could help in selecting true positive epistatic variables more frequently than applying any single method for feature selection via simulation studies. We were able to demonstrate the effectiveness of collective feature selection along with a comparison of many methods in our simulation analysis. We also applied our method to identify non-linear networks associated with obesity.

## Background

The advancements and cost-effectiveness of genotyping and sequencing technologies have led to the ever increasing “short fat data” problem (where the number of features outnumbers the sample size; p>>n) in applying various machine learning methods to detect epistasis[1,2]. Gene-gene interactions are considered as crucial components in the origination of the “missing heritability” for testing association of variants with single or multiple disease traits[3,4]. Various statistical and biological filtering techniques are commonly applied to select variants that are most meaningful in the search for epistatic interactions linked with common and complex diseases[5,6]. Regression approaches are frequently used to model pairwise interactions but many machine learning approaches such as multi-factor dimensionality reduction (MDR)[7,8], neural networks[9], support vector machines[10], Bayesian methods[11], among others are contemporary methods now more commonly applied. Most of these methods are limited in the number of features they can handle, and thus dealing with the computational burden poses a challenge in the application of these methods. Beside the computational burden, it is also important to note that the efficiency of most learning-based methods can be improved to a greater extent if the number of input variables can be reduced. In order to do so, many feature selection methods have been proposed in the past and have been applied in the context of detecting statistical epistasis to identify non-linear associations of genetic variants with a disease trait. Hence, feature selection is not a new concept. Several parametric and non-parametric methods such as LASSO[12], Elastic Net[13], Random Forests[14], ReliefF[15], Gradient Boosting[16], etc., have been developed and used frequently to perform feature selection. All methods have some advantages and disadvantages, and thus they do not follow a “one method fits all” criterion.

In this study, we tested an eclectic set of parametric and non-parametric methods on simulated datasets to first pick a few orthogonal methods to use in selecting features that can be used in downstream analysis of epistasis. We compared these methods based on both efficiency and effectiveness. We observed that different methods tend to select variables based on different important aspects. Thus, we suggest a collective feature selection approach. We propose to select the union of features from the top comparable methods. The concept of taking the input from many algorithms, to select variables as a collective opinion is in line with the “no free lunch” theorem of optimization which states that in searching for candidate solutions, no one algorithm can be specialized to all problems[17]. Unknown genetic etiology of complex diseases makes it theoretically impossible for one algorithm to be specialized in identifying all possible combinations of predictors associated with a disease. This concept is also similar to the concept of “Crowd Machine” which has been explained in previous work[18]. Crowd Machine learning refers to combining multiple machine learning methods into a single machine learning method so that the features from all methods can be used effectively. Our proposed method is a variation of this concept. We recommend applying a collective approach using various top performing feature selection methods to identify variants with varying effect sizes (high and low penetrance) and MAF.

## Methods

In this section, we will describe the datasets used for simulations and real data analyses as well as the statistical methods applied for conducting feature selection.

### Simulation Studies

#### Simulated Data Experiment 1

We simulated multiple data sets consisting of SNPs (single nucleotide polymorphisms), referred to as variables, using an additive genetic encoding (AA=0, Aa=1, aa=2) with case-control status to test for binary outcome. Our simulation parameters consisted of various combinations of the following epidemiological characteristics:

- Disease penetrance: This refers to the strength of the simulated signal or effect size and thus directly corresponds to the heritability of the phenotype. We have used previously simulated data with the same signal strength, these are listed as 0.1_diagonal, 0.5_diagonal and 0.9_diagonal and have been previously explained in Li et al[11].
- Number of disease sites: This refers to the number of SNPs that contributes to the total effect in the dataset.
- Minor Allele Frequency (MAF): For many genetic interaction studies, it has been shown that MAF highly influences power to detect true interactions. Therefore, we limited our analysis to only common alleles above MAF 0.4[19,20]. For main effect variants, we limited the MAF of the causal SNP to 0.4. For interacting effects, MAF for each of the two interacting SNPs was also set to 0.4.
- Number of Samples: We generated 8 simulation scenarios consisting of balanced datasets with 2,000 cases and 2,000 controls.
- Number of Variants: We set the number of variants as 100 and 500 to address the computation burden.

We simulated (a) main effect only and (b) interaction effect only datasets using a simulation procedure that has been previously explained[11]. As described, we evaluated feature selection methods on similar datasets. **Table 1** lists details of all parameters used in generating main effect and interaction effect datasets. For all the simulation analyses, we generated 10 data sets for each combination of parameters in this experiment since we were interested in obtaining mean accuracy values or scores from the replicates to compare across different methods.

**Table 1:** Parameters used for generating simulated experiment 1 data. All datasets consisted of 2,000 cases and 2,000 controls (4,000 samples in total). ‘G’ here refers to the SNP ID prefix.

| Type of Effect<br>(100 and 500 SNPs) | Dataset Name | Causal SNP Model<br>(Penetrance: 0.1, 0.5 and 0.9) |
| --- | --- | --- |
| Main Effect | 1SNP | G1 |
|  | 2SNP | G1, G2 |
|  | 3SNP | G1, G2, G3 |
|  | 4SNP | G1, G2, G3, G4 |
| Interaction Effect | case1_control0 | G1<->G2 |
|  | case1_control1 | G1<->G2<br>G99 <->G100 |
|  | case2_control0 | G1<->G2<br>G3 <->G4 |
|  | case2_control2 | G1<->G2<br>G3 <->G4<br>G97<->G98<br>G99<->G100 |

Using marginal association as an example, the association of a variable with a binary outcome simply indicates differences of allele frequencies between cases and controls. Using a really simple dummy example in **Table 2**, we can see that there are more 1s and 2s for SNP1 in the controls than cases. Thus, in simulation, we could simulate case and control data separately by specifying different allele frequencies for SNP1. Using frequencies as probabilities, we then used sample with replacement until we reached the desired samples.

**Table 2:** Dummy example representing the simulation criteria for main effects in simulation experiment &1. Here 0 in column1 refers to controls and 1 refers to cases. In column 2, 0,1 and 2 refers to the genotypes.

| Case control status | SNP1 |
| --- | --- |
| 0 | 1 |
| 0 | 2 |
| 0 | 2 |
| 0 | 0 |
| 1 | 0 |
| 1 | 0 |
| 1 | 0 |
| 1 | 1 |

The same logic extends to interacting variables. Only in this case, we are specifying different joint allele frequencies between two SNPs. For example, there are more SNP1 = 2 and SNP2 = 2 combination for controls than cases in **Table 3**. We used the different joint allele frequencies to simulate interaction case and control data.

**Table 3:** Dummy example representing the simulation criteria for interacting effects in simulation experiment &1. Here 0 in column1 refers to controls and 1 refers to cases. In column 2 and 3, 0,1 and 2 refers to the genotypes.

| Case control status | SNP1 | SNP2 |
| --- | --- | --- |
| 0 | 2 | 2 |
| 0 | 2 | 2 |
| 0 | 2 | 2 |
| 0 | 0 | 0 |
| 1 | 0 | 0 |
| 1 | 0 | 0 |
| 1 | 0 | 0 |
| 1 | 1 | 1 |

Therefore, simulated variables with interaction effects were generated in case and control datasets separately. For example, in dataset 1SNP, there is 1 SNP with main effect and similarly for interaction effect datasets such as case2_control2, there are 2 pairs of interaction SNPs simulated in cases (G1<->G2 and G3<->G4) and 2 interaction pairs in controls (G97<->G98 and G99<->G100).

#### Simulated Data Experiment 2

We simulated another set of datasets with slightly different architecture to recapitulate the need to consider the varying architectures of complex disease traits. For this simulation, we used GAMETES software to simulate two different genetic architectures[21]. The two architectures here are reflected by the ease of detection measure (EDM) which makes the model easier (EDM-2) or harder (EDM-1) to detect[22]. We simulated 100 features with MAF 0.2 for all predictive features in each dataset. In this scenario, we simulated datasets with a sample size of 2000 (1000 cases and 1000 controls). For each combination of parameters, we simulated 50 replicates at a heritability value of 0.1, 0.2, or 0.4. There were 3 predictive features and 97 non-predictive features simulated in each dataset. The predictive features are included in such a way that there is a pairwise pure epistatic interaction as well as a third main effect additively combined with the interaction.

### Biological Data Application

#### Natural Biological Data

We applied the proposed collective feature selection to a real dataset obtained from the Geisinger MyCode DiscovEHR collaboration[23,24]. At the time of these analyses, the DiscovEHR study consisted of 60,000 samples whose genotype data (using Illumina Human Omni Express Exome chip) is linked to their Electronic Health Record (EHR). For our analysis, we extracted unrelated European American samples of age 18 or older. We extracted all available Body Mass Index (BMI) values for all samples who also had genotype data, from the Geisinger EHR. Median BMI was calculated for all samples and used as the basis for the obesity phenotype in the subsequent analyses. Average BMI of DiscovEHR population is 30[24]. After quality control, 40,449 samples were divided into cases and controls where samples with BMI ranging from 18-24.9 (defined as normal range) were considered as obesity controls and samples with BMI>30 (defined as obese) were considered as obesity cases. We excluded samples in marginal BMI range (25-30) to remove phenotypic heterogeneity and classify samples as normal and obese (extremes of the distribution). To conduct a two-step analysis for feature selection and model testing, we divided the dataset randomly into two parts: variable selection dataset and modelling dataset. Our variable selection dataset was used for feature selection and consisted of 15,201 samples (3,917 controls and 11,284 cases) while our modelling dataset, which contained 14,925 samples in total (3,767 controls 11,158 cases), was used for downstream analyses. We also performed quality control on genotype data to only include variants with genotyping call rate >99%, MAF>20% and HWE P-value<1e-07. Lower frequency variants (MAF<0.2) were excluded from analyses as a first filtration step so as to compare our methodology to the simulated datasets. Additionally, studies also suggest that for variants with MAF <0.2, the interaction effects do not explain much of genetic variance[19,20]. To reduce the search space for testing, we LD-pruned the data to only include independent variants. We used an R^2^ threshold of 0.2 for LD pruning. After genotype QC, the training dataset consisted of 60,232 variants for feature selection.

### Statistical Methods

To compare and contrast the different methods that can be used to select features with non-additive effects, we chose a wide range of filter and embedded methods. For filter-based feature selection methods we tested MultiSURF^*^ and MDR and for embedded methods we tested Random forests, gradient boosting, LASSO and Elastic Net. In this manuscript, we divided methods into parametric and non-parametric methods. We used this terminology throughout the manuscript to classify the methods tested. **Figure 1** lists the three categories of methods we tested for feature selection. We will describe in detail how we ran analyses using these methods in this section. Datasets that are imputed or obtained from commercial genotyping chips such as Illumina consist of 500K to approximately 10M variants. After quality control and LD pruning to include only the most independent variables, it is common to still be left with over 50,000 variants that can be exhaustively tested for interactions. Feature selection can reduce the number of variants and consequently, the computational burden for downstream analysis. Since we chose different methods to test in our analysis, it is important to note that the format of output from all these methods varies and has limited the way in which we can compare the accuracy of these methods. For example, some methods provide a test-statistic for every model, where others provide a ranked list of variables based on performance. In comparing all methods, we could not choose an arbitrary test statistic threshold for each method as that could create bias in selecting variables based on different test statistics as followed in each method (for example MDR and uses balanced accuracy for ranking models, LASSO and elastic net uses lambda for ranking variables, Ranger and gradient boosting uses variable importance measure based on prediction accuracy for ranking variables, etc.). Therefore, to compare all different feature selection approaches, we employed a ranking based method to extract the highest ranked features from a user-defined percentage after running each algorithm. Ranking here refers to the score or the accuracy estimates from each algorithm separately. For all our simulation tests, we showed results for selecting variables at several different user-defined thresholds: 2% or 3% (based on number of effect SNPs in the two sets of simulated datasets), 5% and 10%, thus we select the top 2/3%, 5%, or 10% of the variables in both simulated data experiments to investigate whether the methods perform better or worse, i.e. selection of true positives in comparison to false positives (see **Figure 2**). Next, we ran all of the methods on all replicates of datasets generated from the combinations of parameters as explained in the simulated data section to reduce the error and increase robustness of the models selected. We then averaged across the replicated runs to compare the results.

**Figure 1:**
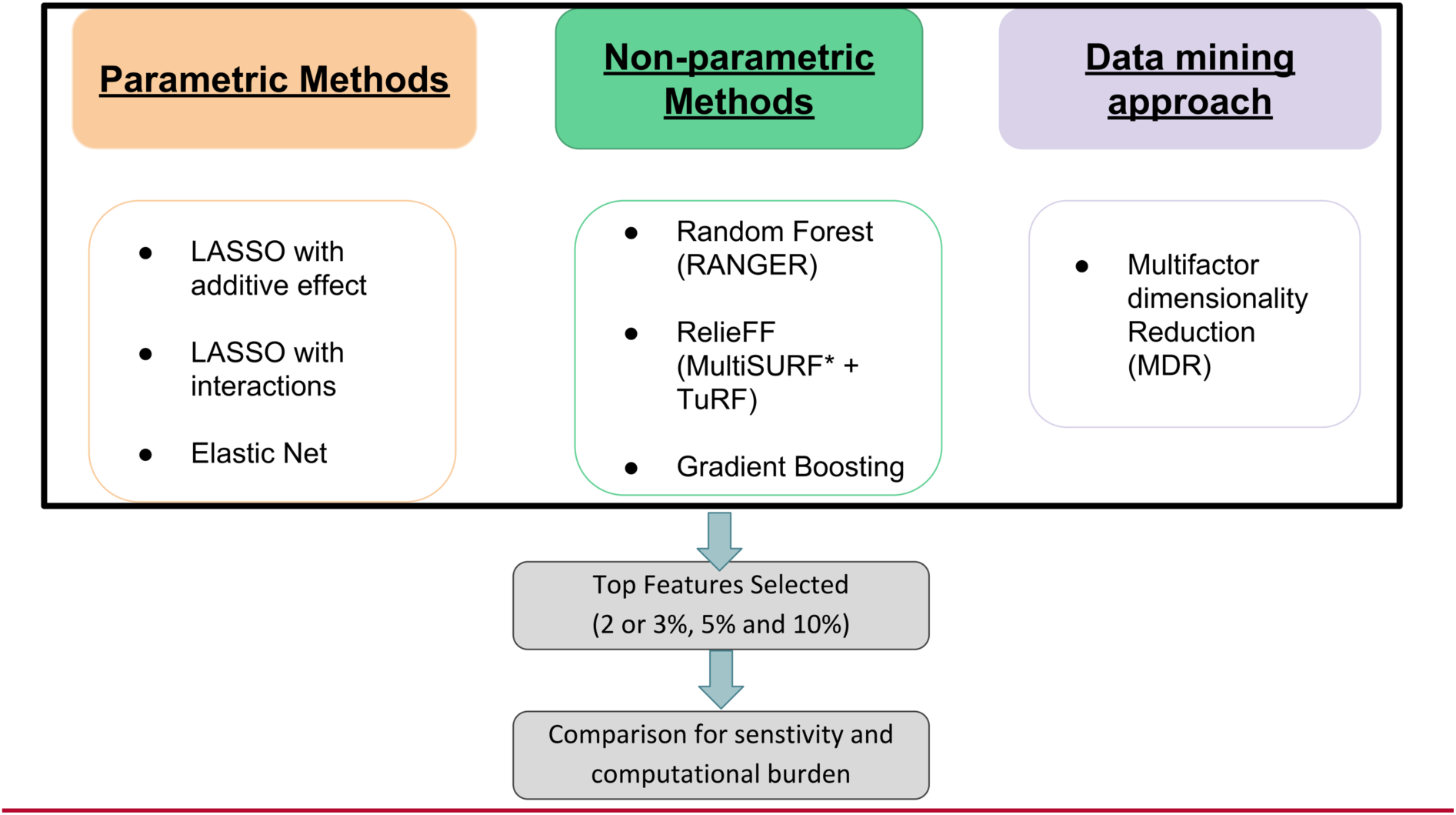
Methods explored for feature selection and selection of top user defined percentage of features for comparison

**Figure 2:**
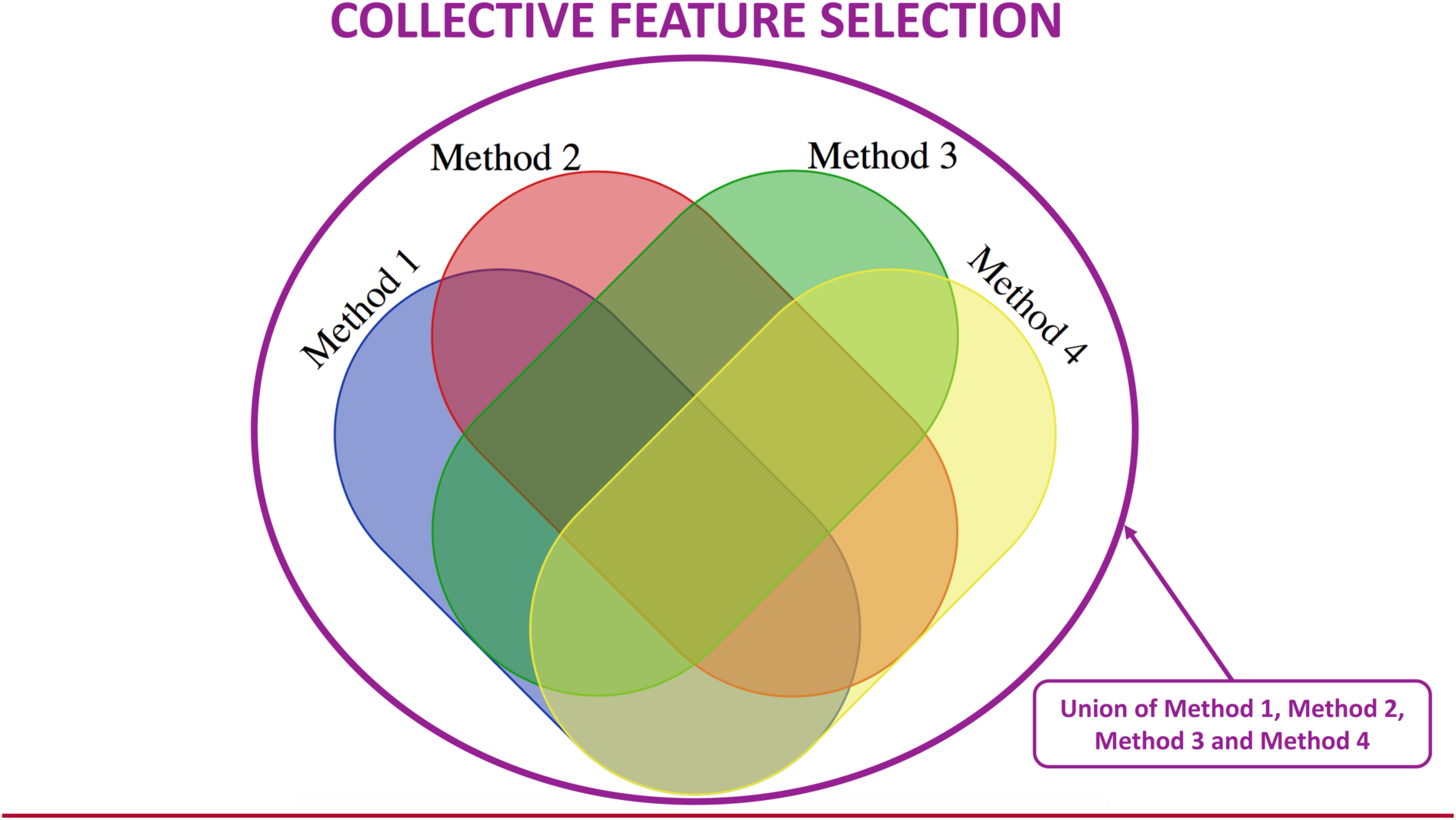
Outer circle represent a collective feature selection approach

#### Parametric Methods

LASSO and Elastic Net regularizations are widely accepted methods for feature selection[6]. Least Absolute Shrinkage and Selection Operator, or LASSO[12,25], is a shrinkage and variable selection method with imposed *L1* regularization on the regression coefficients. Since the main goal of this analysis is to detect features that exhibit interaction effects, we ran LASSO regression to include both additive effects of SNPs in the models and exhaustive pairwise interactions of all SNPs in the model. Below are the equations representing LASSO regularization for single SNP and interactions:

Penalized estimates for the model with additive effects alone can be derived as the solution to the following optimization problem:

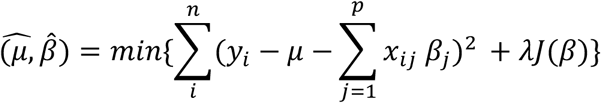

where 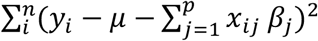 is the residual sum of squares, λ >= 0 is the regularization parameter and *J*(*β*) is the penalty function.

For LASSO Regularization with additive effects only, the penalty function is the L1 norm and can be expressed as follows:

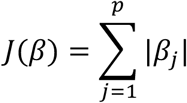

Likewise, penalized estimates for the regression model with additive and SNP-SNP interaction effects can be derived as the solution to the following optimization problem:

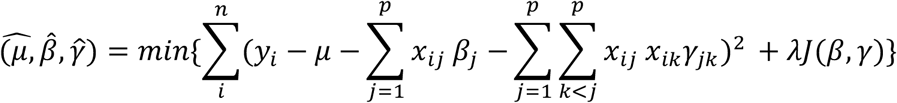

Again, for LASSO Regularization with additive effects and interactions together, the penalty function can be expressed as follows:

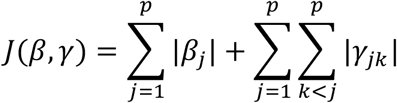

LASSO combines variable selection and shrinkage of variables, but it has a drawback when the number of predictors is greater than the number of samples (p>n), in which case it tends to select at most n predictors. Also, when predictors are correlated, LASSO is outperformed by ridge regression. Thus, we modeled the data with ridge regression in a preliminary part of our analysis but did not include those results in this manuscript since they were similar to those from LASSO. Next, we explored another penalized regression method, the Elastic Net, which works well in selecting a group of correlated variables and does not limit the selection of the number of variables. Elastic Net uses a weighted average of the *L1* and *L2* norms for its penalty function.

Similar to the LASSO penalty function, the elastic net penalty function [Zou et al; 2005] for the model with additive effects and interactions can be expressed as follows:

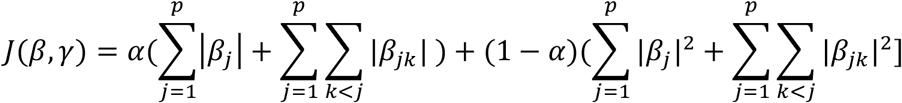

Both these penalized regression methods (LASSO and elastic net) require optimization of 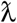. Elastic net involves another tuning parameter called *α*, which is commonly set to 0.5. In order to help tune these parameters, we performed 5-fold cross validations for these two methods and chose the most optimal regularization parameter for feature selection.

#### Non-Parametric Methods

Even though parametric methods are simple and easy to understand, they do not always fit the complex nature of biology. Thus, exploring some non-parametric methods is also necessary. Non-parametric methods do not make assumptions about the distribution of variables and underlying genetic architecture. These methods usually work best for “big data” problems. We tested two decision-tree based methods, including Random forests and Gradient Boosting, and we also tested a non-heuristic ReliefF algorithm variation called Multiple Threshold Spatially Uniform ReliefF (MultiSURF^*^)[26].

For our random forests implementation, we used the RANGER R package[27]. We tuned random forests to get better results, setting number of trees as 1,000 for main effect datasets and 4,500 for datasets with interaction effects. The other parameter that we tuned is the number of variables that split each node; we used 35 for main effects, 70 for interaction effects in datasets with 100 SNPs, and 200 for interaction effects in datasets with 500 SNPs. We also used gradient boosting implementation in the GBM R package. For gradient boosting, we set the number of trees as 800 for main effect datasets and 15,000 trees for interaction effects datasets. We set the bag fraction as 0.5 and shrinkage as 0.01, which have been suggested to result in the best performance based on the best practices from R package manual (https://cran.r-project.org/web/packages/gbm/gbm.pdf). TuRF refers to Tuned ReliefF and it performs feature selection recursively. It is suggested to use TuRF along with ReliefF algorithms to get better performance when using a large number of variables[5,15,28].Thus, it is important to test the number of variables that will be thrown out at every iteration. We tested discarding 1%, 5% and 10% of least predictive variables at each iteration to determine the appropriate threshold for MultiSURF^*^+TuRF in order to identify more true positives in a computationally feasible amount of time.

#### Non-parametric data mining approach for feature selection

Multifactor Dimensionality Reduction (MDR) has been traditionally applied to several association studies including gene-gene and gene-environment interaction studies[7,8]. Many different versions of MDR have also been proposed for different data types[29–31]. Using MDR as filtering method has also been previously tested and compared with other methods[32]. We utilized parallel MDR (pMDR) {https://ritchielab.psu.edu/software/mdr-download} in similar way to Oki NO et al[33], where we ran all main effect models and two-way interactions without cross validations and then ranked the variables based on their training accuracy.

#### Collective feature selection

A plethora of machine learning and feature selection methods have been proposed and tested in various studies[6,12,13,16,32]. In this manuscript, we aimed to compare a few of these methods; however, picking one method can be convenient but not always pertinent. Thus, in our analysis we proposed to select a few orthogonal feature selection methods from what were tested and then use the union of all variables selected from these methods for any downstream analysis. **Figure 2** depicts the concept of our collective feature selection approach in selecting variables in a dummy set of 4 methods listed as Method 1 to 4. We applied this approach to simulated data experiments 1 and 2 but are only showing results from experiment 2 where we selected top 3%, 5% and 10% features from each of the 4 methods. Results from experiment 1 are similar. We compared these methods in terms of number of overlapping features, number of true positives, and number of false positives detected from each. To represent the overlapping features and features selected for EDM-1 and EDM-2 models, we merged the datasets with 3 different heritability values (0.1,0.2 and 0.4) together.

Using this approach, we also propose a pipeline as shown in **Figure 3** for performing analysis using feature selection as an essential step before applying machine learning methods, such as neural networks, support vector machines, Bayesian approaches, etc., in downstream analyses. Figure 3 represents a three-step pipeline, beginning with testing several feature selection methods in simulated datasets in Step 1, as covered in this manuscript. Steps 2 and 3 involve applying the selected methods to a real dataset. We propose to apply top performing methods from Step 1 on our natural biological dataset in Step 2. In Step 2, we select variables based on our collective approach in our real training dataset. Finally, in Step 3, we propose to extract collectively selected variables from the variable selection subset of our natural biological dataset to then use for downstream analysis.

**Figure 3:**
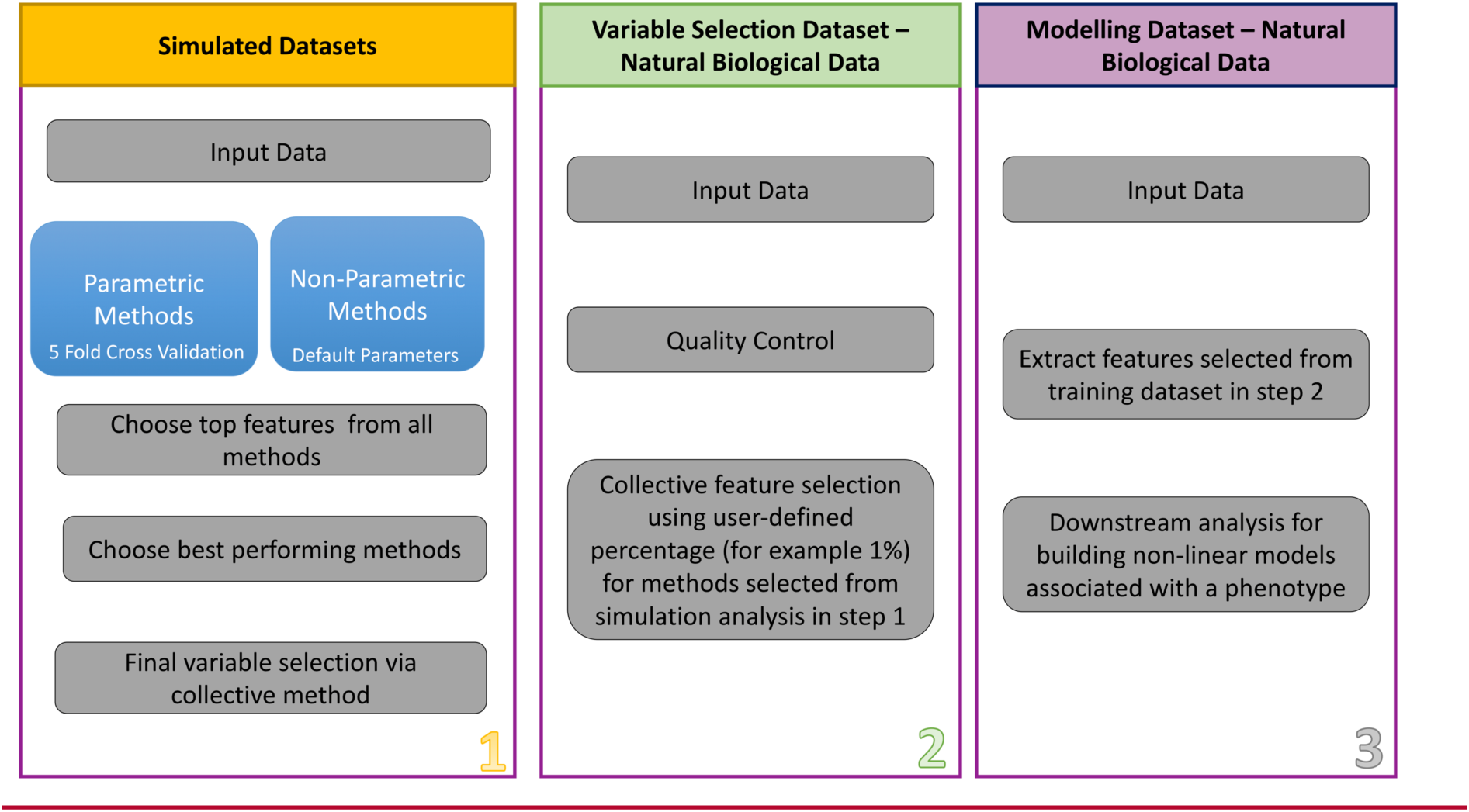
Pipeline of feature selection procedure and downstream analysis in both simulated and natural biological data.

#### Feature Selection and Downstream Analyses

We applied this proposed approach to test for SNPs that are associated with obesity among samples from the MyCode DiscovEHR study[23]. On quality controlled data, we selected features using MDR, MultiSURF^*^ and TuRF, and Ranger collectively, and then performed downstream analyses using Analysis Tool for Heritable and Environmental Network Associations (ATHENA)[9,34]. We choose to apply Grammatical Evolution Neural Networks (GENN) implemented in ATHENA for this analysis to select non-linear epistatic interactions between SNPs selected from the feature selection strategy described above. Grammatical evolution methods are alternatives to classical genetic programming approaches in machine learning methods. This approach has been widely accepted and its effectiveness has been explained in previous studies[35–38]. We used the following parameter criteria to identify networks associated with BMI case control outcome:

1. 5-fold cross validation: Modelling data as described in Step 3, which included 14,925 samples and features selected via collective approach, were divided into 5 equal parts
2. Process: The first iteration begins with selecting a training set to generate random population (popsize 10,000), dividing into sub-populations, and then preforming an analysis on 30 nodes. The grammar for GENN is then used to evaluate the training set using Area Under Curve (AUC) fitness criteria. This step is then repeated 20 times (numsteps) after which migration takes place to select the best solution from all 30 nodes. This process is repeated 4 more times, once for each remaining cross-validation fold to perform 5-fold cross validation as explained in step #1.
3. Results: Training and testing AUC for each network model associated with outcome is reported from all cross validations.

## Results

### Simulation Studies Results

#### Optimizing TuRF iterations

We aimed to use TuRF along with MultiSURF to help increase its efficiency. We tested 3 different thresholds, 1%, 5%, and 10%, to iteratively remove that percent of lowest ranking variables at each iteration. **Figure 4** below shows the comparison of results. Here sensitivity is defined as the proportion of true positives selected where a sensitivity of 1 means 100% of true positives were selected from the simulated dataset.

**Figure 4:**
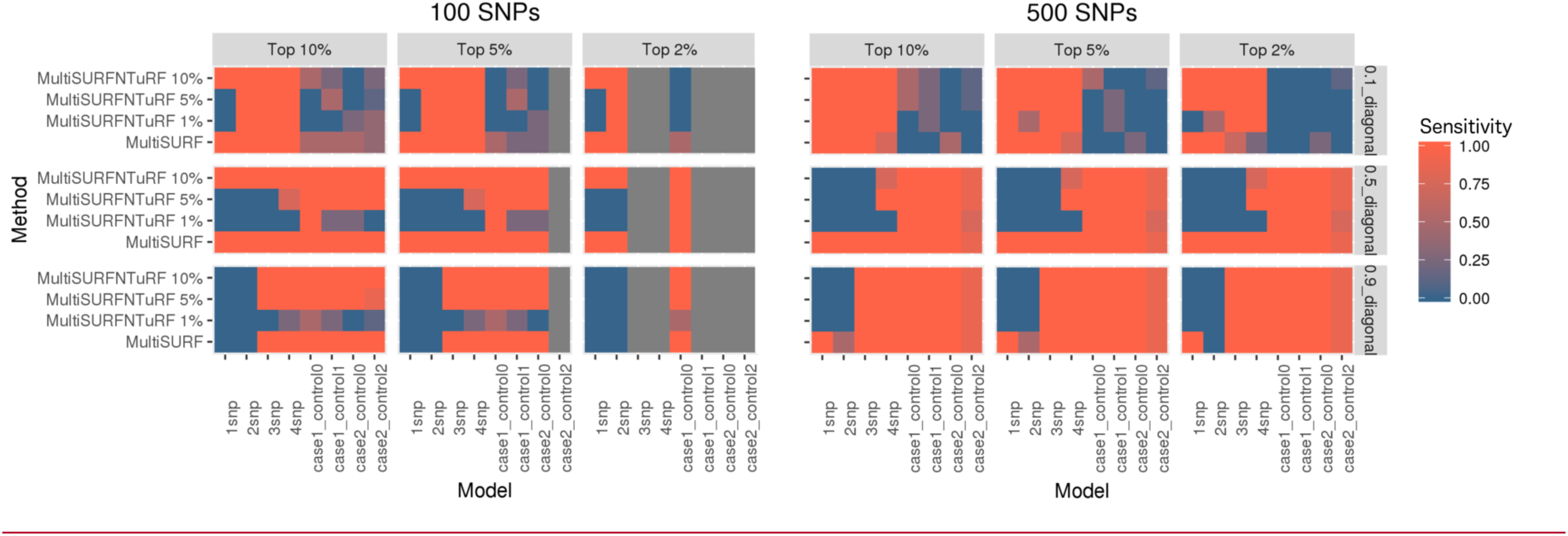
Comparison of results for TuRF parameters when using it with MultiSURF^*^ as well as MultiSURF^*^ without TuRF implementation. The plot on left is for 100 variables and the right plot is for 500 variables. The x-axis lists main effect and interaction datasets while the y-axis lists all methods tested. These plots are faceted by percentage of top variables selected and the strength of signal. The color gradient refers to the sensitivity (percentage of true positives), ranging blue to orange, or 0 to1.

It is interesting to note here that MultiSURF^*^ without TuRF performs better for strong effect models (0.5 and 0.9 penetrance) for both 100 SNPs and 500 SNPs datasets. This could be due to the fact that TuRF works better for larger datasets with many variables whereas 500 variables is still considered relatively “small” and can be handled by MultiSURF alone. In this case, TuRF does not help but instead makes it worse. The poor performance of MultiSURF and TuRF could be explained by the algorithm accidentally discarding the important variable or variables in the first iteration.

#### Distribution of accuracy from MDR

We ranked all MDR generated models based on their training accuracy to select top user-defined percentages of models as explained in model selection. **Figure 5** shows the distribution of median training accuracy for all models that were selected from MDR feature selection. It is to be noted that the accuracies for the selected features vary greatly based on the strength of signal. For example, training accuracies in 0.1 effect signal datasets are close to 55% for all models whereas accuracies for 0.9 effect signal datasets are closer to 80%. Notably, in many two-way interaction models, we observed that false positives are paired with true positives. Figure 5 represents overall accuracy of the model. Since both false and true positives exist in model, the accuracies reported are also higher for false positive (in red).

**Figure 5:**
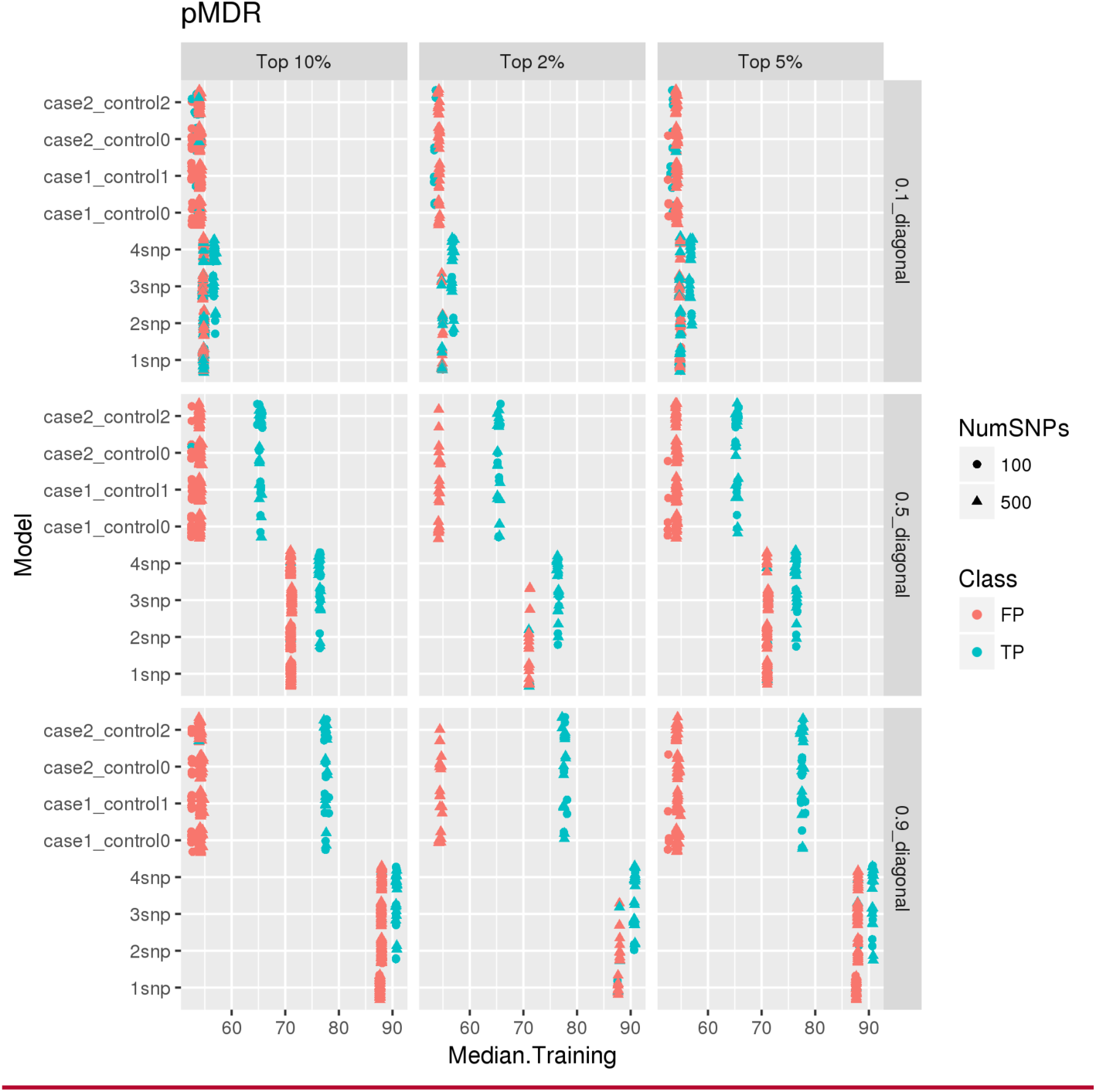
Distribution of median training accuracy from pMDR analyses. X-axis is median training accuracy values of the model, Y-axis lists all main effect and interaction simulated datasets. These plots are faceted by effect size and top percentage of models selected. 100 SNP data is shown in circles and 500 SNP in triangles. The two colors represent actual true and false positives in results.

#### Application of all methods on simulated datasets

We tested all chosen methods on the two experiments of simulated datasets with different ranges for effect sizes and various additive main effect and interaction effect models as explained in the data section. To compare results, we are using the degree of effectiveness described as “Sensitivity” where a sensitivity of 1 is equivalent to 100% of true positives being selected in the top features. **Figure 6** represents the results for all methods tested using simulated data experiment 1 and **Figure 7** represents results from all methods tested using simulated data experiment 2. It seems evident from these plots that MDR used as a feature selection tool helps to select true positives every time for models tested in simulated data experiment 1 whereas Ranger and Gradient Boosting perform best in terms of selecting true positives for data experiment 2. In the first set of simulations, we see that nonparametric methods do not perform as well in detecting interacting effects. Additionally, we see that most methods do not perform well for the weakest signal tested (0.1 penetrance). MultiSURF and TuRF seem to perform well for interaction effects but do not perform well for main effects. In the second set of simulations, LASSO (without explicitly adding interactions in the model) and elastic net fail to find true positives in both EDM-1 and EDM-2 models, while MDR fails to identify true positives for EDM-2 model. MultiSURF alone and MultiSURF with TuRF both struggle in finding true positives from EDM-2. MDR and ReliefF algorithms work well for EDM-1 model architecture. Lastly, LASSO with interaction models can identify interactions similar to best performing methods in both simulated sets. When we compared these methods for their efficiency (computation burden and memory requirements) as shown in **Figure 8 and Table 4**, we observed that the parametric methods (especially LASSO with interactions) take more computation time than most non-parametric methods and the data mining approach. Additionally, LASSO generates an pxp matrix (SNPxSNP) for all exhaustive pairs of SNPs and also requires more memory than other methods to perform computation. Methods like LASSO and Ranger (R package) were also not computationally feasible to run on large genome-wide datasets including over 50,000 SNPs. Thus, a pre-filtration of SNPs based on criteria like LD pruning, MAF filter would be necessary.

**Table 4:** Computational time and memory requirements for all feature selection methods, compared in terms of number of SNPs. Note that “NA” here stands for where the model could not be tested due to computational infeasibility while keeping all parameters for simulated datasets same.

| Method | Computational time based on number of SNPs (in seconds) |  |  |  | Memory requirements based on number of SNPs |  |  |  |
| --- | --- | --- | --- | --- | --- | --- | --- | --- |
|  | 100 | 500 | 50,000 | 100,000 | 100 | 500 | 50,000 | 100,000 |
| LASSO | 4.65 | 58.49 | NA | NA | 10gb | 10gb | NA | NA |
| LASSO with interactions | 186.9 | 1800 | NA | NA | 20gb | 20gb | NA | NA |
| Elastic Net | 31.44 | 401.11 | NA | NA | 10gb | 10gb | NA | NA |
| Ranger | 151.3<br>1 | 681.83 | NA | NA | 8gb | 8gb | NA | NA |
| Gradient Boosting | 103.2<br>2 | 466.95 | NA | NA | 8gb | 8gb | NA | NA |
| MDR | 0.25 | 15.03 | 6102 | 89777 | 1gb | 1gb | 10gb | 30gb |
| MultiSURF | 2.48 | 5.13 | NA | NA | 18gb | 39gb | NA | NA |
| MultiSURF + TuRF 0.05 | 36.72 | 65.72 | 4420 | 8321 | 18gb | 39gb | 28gb | 28gb |

**Figure 6:**
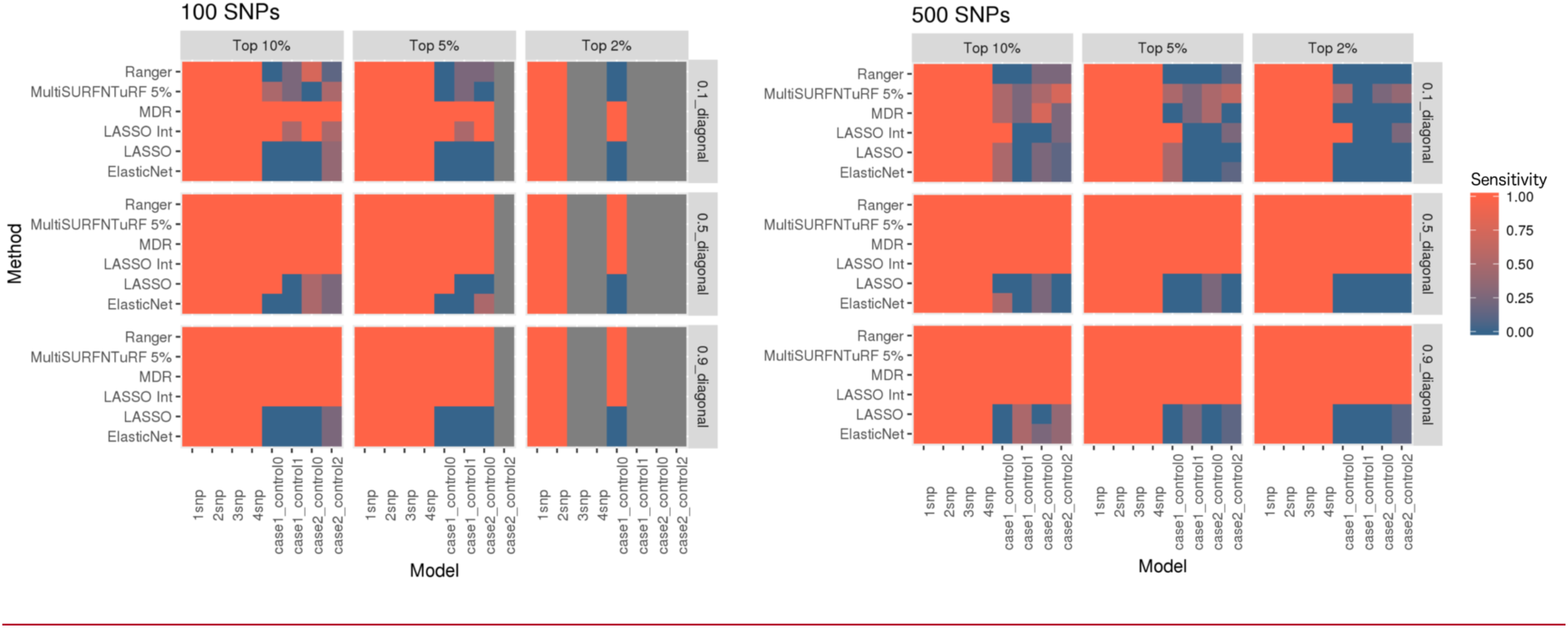
Comparison of results from all methods tested on simulated dataset 1. These heat maps show the sensitivity of results for all methods (on y-axis) and all simulated models (on x-axis) for both 100 SNPs and 500 SNPs datasets in combination with different effect sizes and selection percentage of top features.

**Figure 7:**
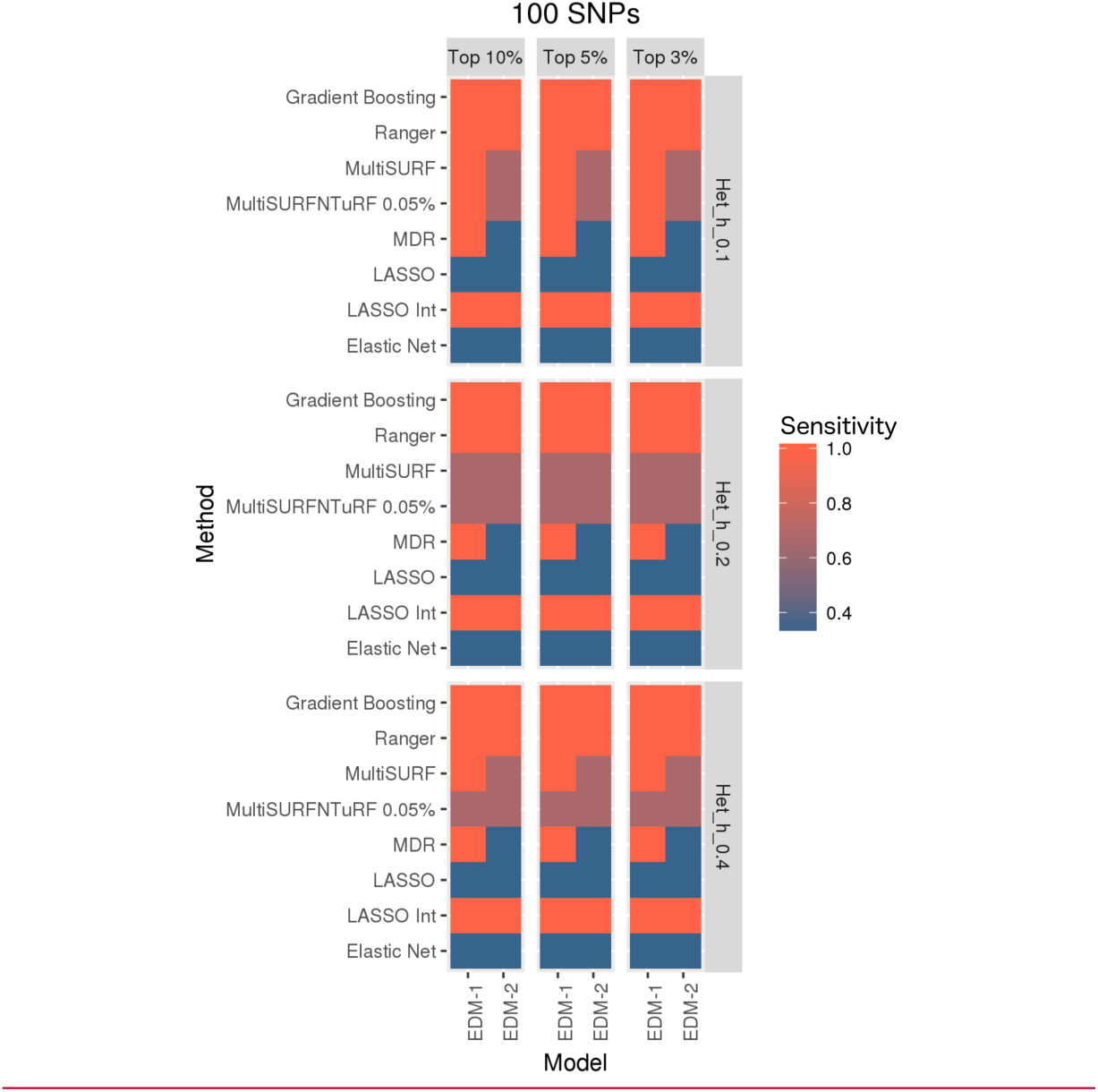
Comparison of results from all methods tested on simulated dataset 2. These heat maps show the sensitivity of results for all methods (on y-axis) and both simulated models (on x-axis) in combination with different effect sizes (heritability values of 0.1, 0.2 and 0.4) and selection percentage of top features.

**Figure 8:**
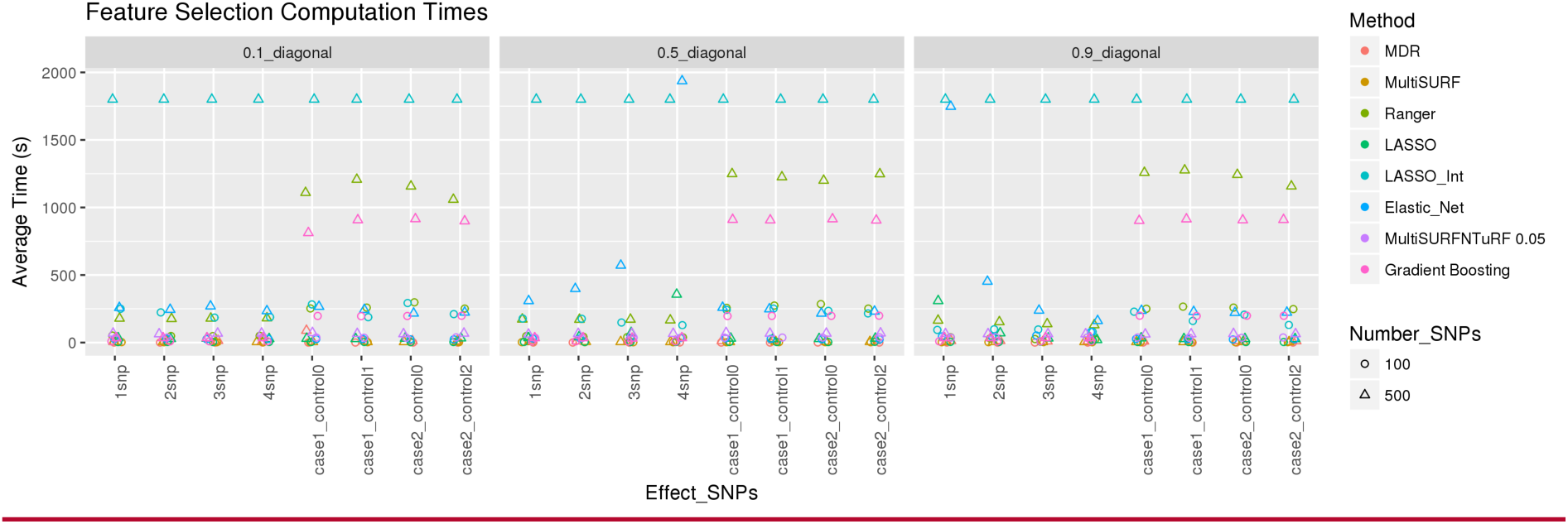
Plot showing time in seconds (on y-axis) taken for running all feature selection methods. All simulated models are presented in x-axis. Color represents each method. Circles are for 100 SNPs datasets and triangles for 500 SNPs datasets.

We also estimated the time it would take for most of these methods to run when the number of samples and variants are increased. From our analysis, we estimated that MDR scales linearly with number of samples and quadratically with number of features whereas MultiSURF^*^ scales quadratically with number of samples and linearly with number of features.[15]. Thus, the computational burden or MDR increases more when number of samples is increased. Gradient boosting seems to only perform well with larger effects sizes in terms of detecting true positives[16] and fewer variants. For larger numbers of variants and samples, the best way to perform analyses using gradient boosting is to create subsets of SNPs with low intercorrelations and then aggregate results (email conversation with Dr. Gitta Lubke). Since we do not want to make any pre-assumption about the nature of interactions and only test subset of SNPs in a smaller region of genome, we decided to not use Gradient Boosting in such manner.

#### Collective Feature Selection on Simulated Dataset

We applied collective feature selection on simulated experiment data 2 to obtain the number of features that will be selected from top performing methods. Figure 9A and 9B show the overlap among top features selected from MDR, Ranger, Gradient Boosting, and MultiSURF^*^ and TuRF on EDM-1 and EDM-2 model architectures. Based on information known about merged results from simulated datasets, we expected to obtain 9 true positives (3 from each heritability parameter) in each set of top features selected by every method. However, we again observe that each method does not pick all true positives as shown in 3rd panel of Figure 9.

**Figure 9:**
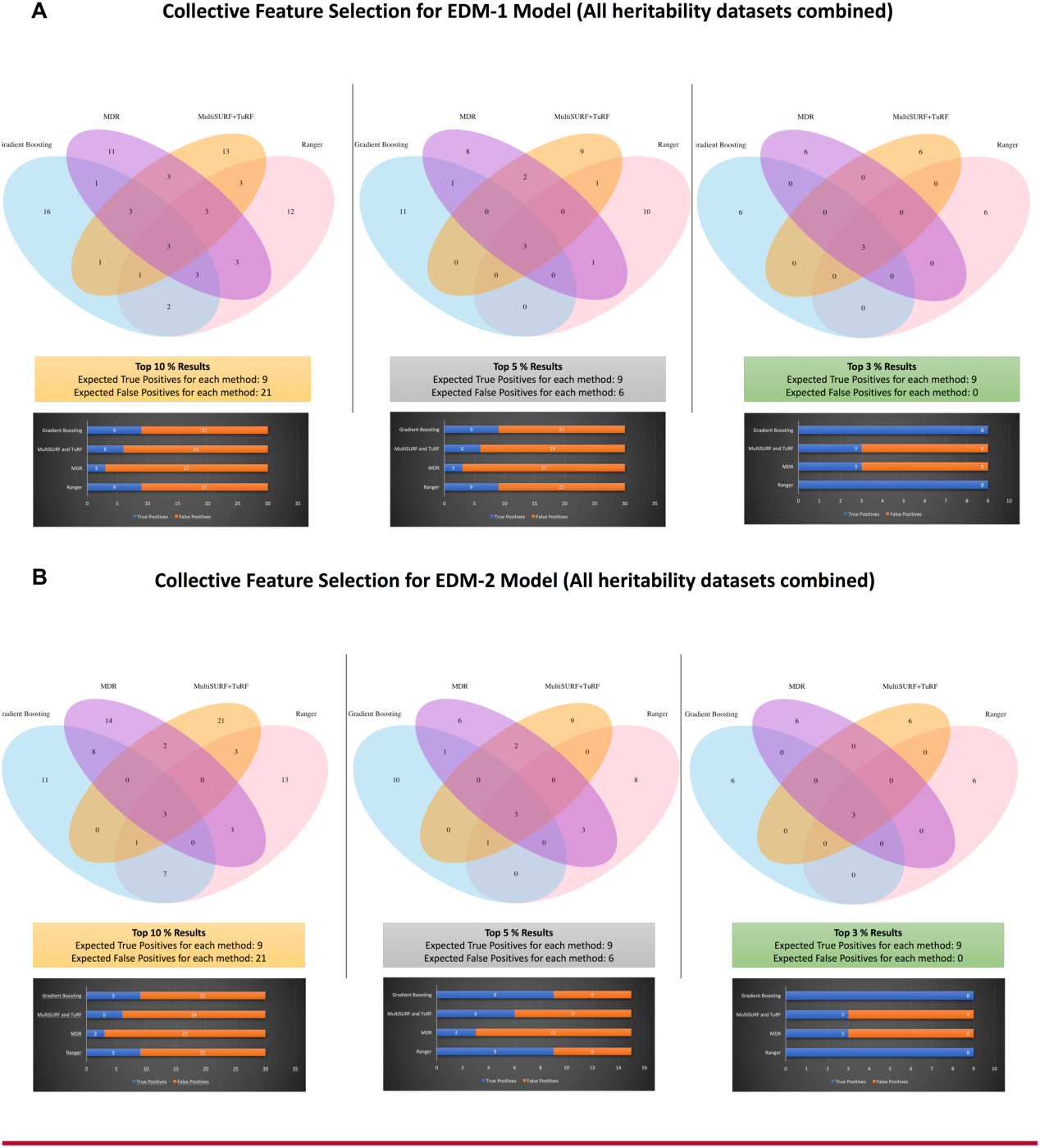
Venn Diagrams shown here represent the overlap among the top features selected by all methods while bar charts below each Venn diagram show the number of true positives and false positives selected by each method. Plot A illustrates results for EDM-1 datasets and Plot B contains results for EDM-2 datasets.

Therefore, the practice of applying a collective approach seems advantageous. Figure 10 shows the number of features selected in each model by top 3, 5, and 10% model selection criteria. One point to note is that by choosing collective feature selection, we picked all 9 true positives every time whereas by picking one method alone, we risk the chance of picking the “best” method based on one scenario and applying it to a dataset where it is unable to detect all of the true positives.

**Figure 10:**
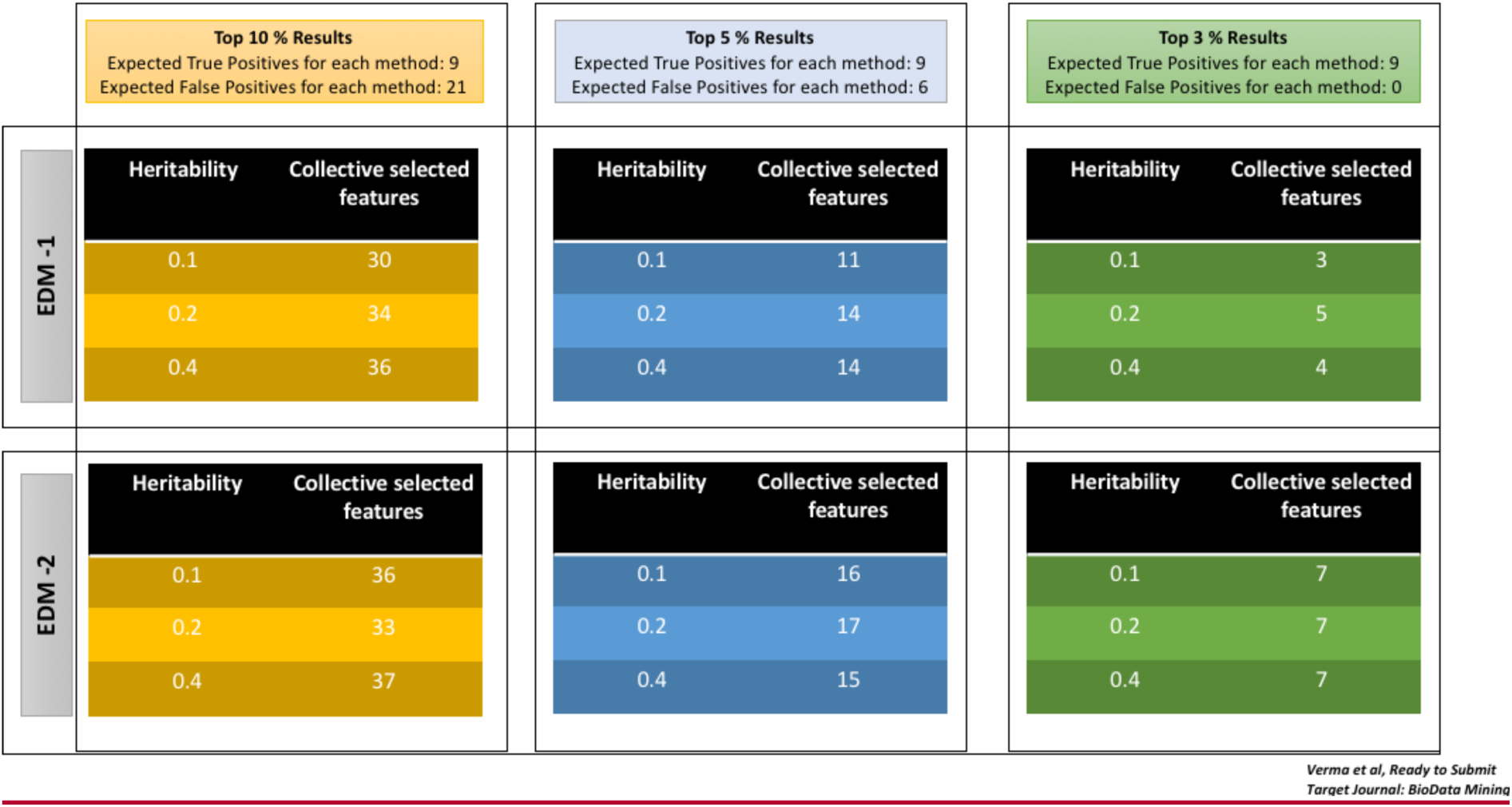
Number of features selected by collective approach

### Biological Data Application

#### Collective Feature Selection

The first step in identifying non-linear models associated with obesity (defined here based on BMI values) is to perform feature selection. We selected 3 methods (MultiSURF and TuRF, MDR and RANGER) for feature selection as described in the methods section. As shown in Table 4, Ranger R package was not computationally feasible (in terms of memory) to run on >50,000 SNPs; we performed feature selection via random forests by combining Ranger with GenABEL R package to load GWAS data. The computational time for collective feature selection is the combination of the time it took to run each method which is 13 days for Ranger + 1 day for MultiSURF and TuRF + 3.5 days for MDR = 17.5 days. Input data consisted of 60,032 SNPs and after feature selection, we selected the top 1% results from each method. This resulted in 1,758 variables selected using collective feature selection (note that intersection of methods only selects 2 genes which do not include well known SNPs linked to obesity such as variants in *FTO* and *MYO16*). The overlap of these variables among the different methods is shown in Figure 11.

**Figure 11.**
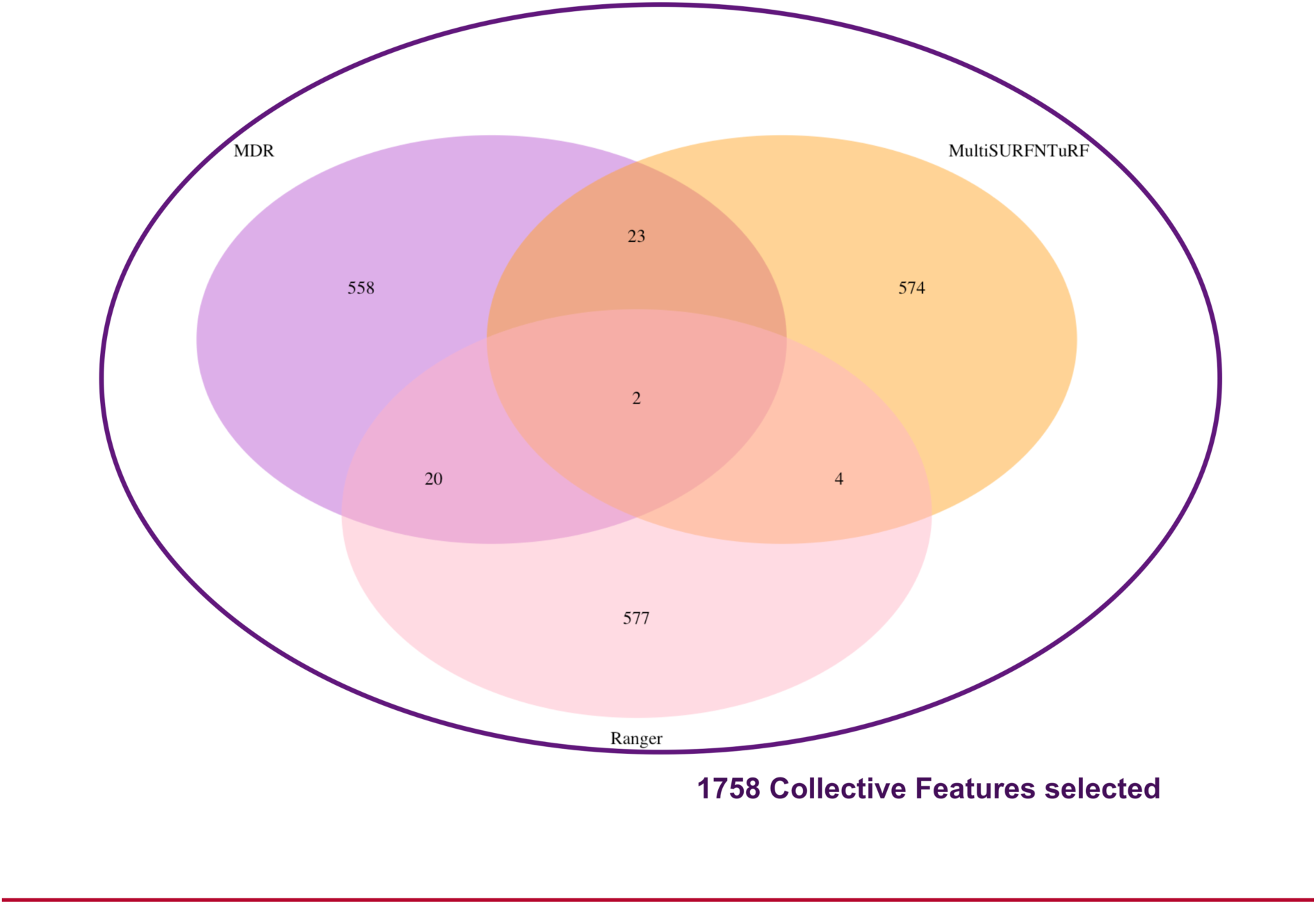
Collective feature selection to select 1758 variables with potential epistatic effect from MyCode data

#### ATHENA Results

5 different networks were obtained as a result of applying GENN to identify non-additive interactions associated with BMI outcome. The training and testing area under curve (AUC) for the 5 models are presented in Table 5.

**Table 5:**
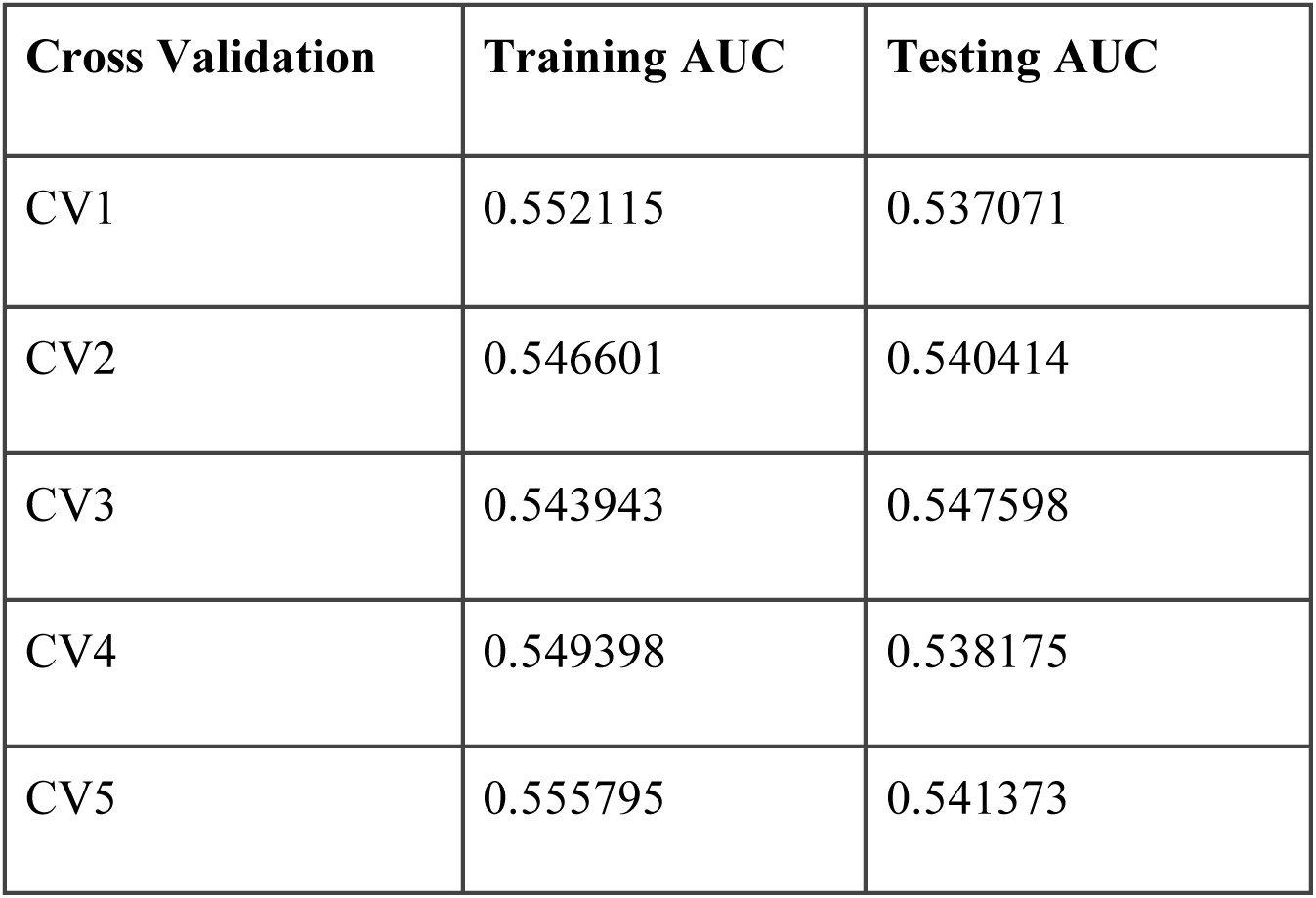
Training and testing AUC for models selected by ATHENA

We choose the best network from this analysis, which is shown in Figure 12. Figure 12 also represents the selection of variants by each feature selection method. In this analysis, we did not adjust for any confounding effects of age, sex, or principal components (PCs) on BMI, but for the variants selected in top models from ATHENA, we ran regression using PLATO[39] to see if the effect sizes and P-values for these variants change drastically when BMI, the dependent variable, is adjusted by covariates (age, sex and first 4 PCs). Therefore, to identify if there is significant effect of co-variates on SNPs, we tested these variants by running logistic regression with and without adjusting for covariates. **Table 6** lists the p-values and betas from regression analyses.

**Table 6:** p-values and betas from regression analyses on 5 SNPs in the network selected by ATHENA

| SNP | Gene Name | No Covariates |  | With Covariates |  |
| --- | --- | --- | --- | --- | --- |
|  |  | p-val | beta | p-val | beta |
| exm-rs11075987 | *FTO* | 6.76E-11 | 0.173939 | 8.02E-09 | 0.1690 |
| rs7232886 | *N/A* | 9.36E-09 | -0.15448 | 6.24E-06 | -0.1336 |
| rs2756184 | *TTBK1* | 0.014104 | -0.06635 | 0.018812 | -0.0699 |
| rs9520911 | *MYO16* | 0.045223 | 0.053358 | 0.051185 | 0.0573 |
| rs7043308 | *TNC* | 0.768626 | -0.00958 | 0.745697 | -0.0116 |

**Figure 12:**
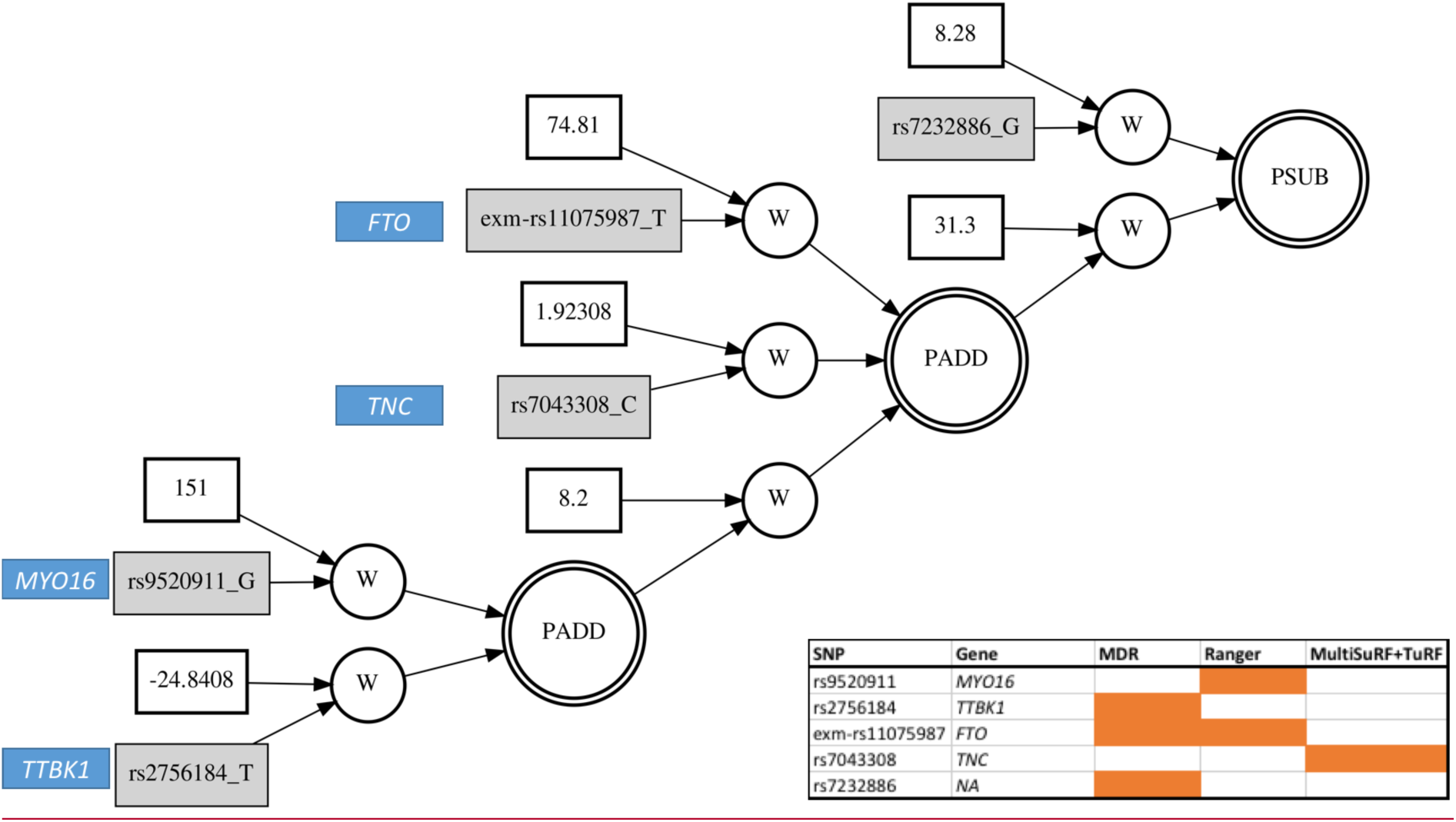
Best GENN model selected from ATHENA. The SNPs are annotated to gene names. On the bottom right is the list of variants and genes in the model and which feature selection method selected the variant are colored in the table to represent the presence (in orange) and absence (in white) of variant in each feature selection method.

Obesity is a worldwide epidemic and it predisposes to many other metabolic traits and diseases[40]. In our network, we observed a well-known hit for a variant in the *FTO* gene which has been identified by many GWAS analyses. It is to be noted that *FTO* variant was not selected by every feature selection method and similar is the case for other variants that are reported in Figure 12. The reported model suggests interaction of *FTO*[40,41] variants with variants in *TNC*, *MYO16*, and *TTBK1* genes. Notably, these three genes have known associations with other phenotypes influenced by BMI, such as *TNC* with Alzheimer’s and schizophrenia[42,43]. *TTBK1* is also known to be associated with Alzheimer’s disease[44–46] while *MYO16* has been found to be associated with pulse pressure[47]. It is also interesting to note that variants in genes *MYO16* and *TNC* are not significant when tested for independent main effect (as reported in **Table 6**) but they are included in the interaction model as suggested by ATHENA (**Figure 12**) which suggests that these variants might work in combination to affect the etiology of obesity but would not be identified otherwise in an additive model.

## Discussion

Epistatic features of genes are necessary to consider when investigating the genetic etiology of disease traits. Gene-gene interactions are believed to account for hidden genetic variability[48]. Testing exhaustive pairwise or higher order interactions among all genetic variants poses various challenges including computational burden and correction for multiple hypotheses. Along with these challenges that affect efficiency, it is also important to note that adding more variables to test also reduces the effectiveness of the predictions. Thus, performing feature selection before modelling is necessary. In our study, we tested parametric, nonparametric, and data mining approaches for feature selection and compared them based on the top models selected as well as the computational time. Through our simulation experiments, we observed that every method is trained to pick variants based on different underlying models that could have potential epistatic effects on disease traits which is reflected by the selection of different false positives from each method on our simulated datasets. Similarly, every method that we tested does not pick all main effect variables every time. This is evident from the non-selection of *FTO* variant by MultiSURF+TuRF and non-selection of variants in genes *MYO16* and *TTBK1* by MDR and Ranger respectively. One possible explanation for selection of different features from different algorithm corresponds to the “no free lunch” theorem[17] and the understanding that no particular feature selection method is specifically designed to pick all epistatic effects. We recommend selecting a user-defined percentage based on combination of sample size, number of variables and trait complexity to obtain the union of features from all methods, referred to here as collective feature selection, to potentially increase power to detect more biologically pertinent associations. It is likely that using a collective approach could result in adding more noise to the analysis, but our analysis suggests that applying different feature selection strategies yield such majorly dissimilar results that the payoff is greater than the cost. In future studies, we aim to test the collective feature selection approach on other natural biological datasets. Our simulation analysis showed that applying non-parametric approaches, like MDR, random forest, gradient boosting and ReliefF results in selecting more true positives epistatic effects in a computationally feasible amount of time than using parametric approaches. But using one method does not always yield all true positives. Thus, we propose collective feature selection utilizing non-parametric methods as a powerful approach for epistatic discovery analysis.

One of the limitations of this study is that we tested our analyses for binary outcome in both simulated and natural datasets. Future work would include the application of these methods to quantitative phenotypes. Additionally, in our simulation analyses we were not able to identify any patterns among the SNPs that were selected across methods. One possible reason could be because of the way that noise is simulated in our dataset, we selected all variants at similar MAF. More studies including simulations of different sets of MAF could also help validate this approach further. In addition, the inclusion of other types of underlying models of epistasis would be useful to further discern which orthogonal or complementary methods perform best in a collective feature selection strategy.

## Conclusions

Although our current study is limited in terms of the simulations we performed, they clearly indicate that different methods select varying features depending on the genetic architecture of the trait. Thus, using a collective approach by selecting union of results from different methods rather than selecting an intersection could help preserve features with non-additive effects during feature selection. We applied our approach to select features that were later tested in an independent dataset to identify networks using GENN. Our model was able to select known signals as well as potential interacting effects of known signals with other variants that could be influencing the risk of obesity.

## List of abbreviations

ATHENA: Analysis Tool for Heritable and Environment Network Analysis
BMI: Body Mass Index
CFS: Collective Feature Selection
GENN: Grammatical Evolution Neural Networks
GWAS: Genome wide Association Study
MAF: Minor Allele Frequency
MDR: Multifactor Dimensionality Reduction
SNP: Single Nucleotide Polymorphism
SURF: Spatially Uniform ReliefF
TuRF: Tuned ReliefF

## Declarations

### Ethics approval and consent to participate

Not Applicable

### Consent for publication

Not Applicable

### Availability of data and material

Additional information for reproducing the results described in the article is available from authors upon request. Availability of natural biological data from DiscovEHR cohort may subject to data user agreement.

### Authors’ contributions

SSV, AML and XYZ performed analyses on simulated and real data. Simulated study data was provided by RL and RU. SSV, YV, DK, MDR were involved in designing the study and appropriate pipelines. SD developed software for running ATHENA and parallel MDR analyses. SSV, DK, YV, MDR participated in drafting and finalizing the manuscript. All authors read and approved the final manuscript.

#### Acknowledgements

We acknowledge discussions at EDGE conference and inspirations from Dr. Jason Moore and Dr. James Malley

